# Human to yeast pathway transplantation: cross-species dissection of the adenine de novo pathway regulatory node

**DOI:** 10.1101/147579

**Authors:** Neta Agmon, Jasmine Temple, Zuojian Tang, Tobias Schraink, Maayan Baron, Jun Chen, Paolo Mita, James A. Martin, Benjamin P. Tu, Itai Yanai, David Fenyö, Jef D. Boeke

## Abstract

Pathway transplantation from one organism to another represents a means to a more complete understanding of a biochemical or regulatory process. The purine biosynthesis pathway, a core metabolic function, was transplanted from human to yeast. We replaced the entire *Saccharomyces cerevisiae* adenine de novo pathway with the cognate human pathway components. A yeast strain was “humanized” for the full pathway by deleting all relevant yeast genes completely and then providing the human pathway in trans using a neochromosome expressing the human protein coding regions under the transcriptional control of their cognate yeast promoters and terminators. The “humanized” yeast strain grows in the absence of adenine, indicating complementation of the yeast pathway by the full set of human proteins. While the strain with the neochromosome is indeed prototrophic, it grows slowly in the absence of adenine. Dissection of the phenotype revealed that the human ortholog of *ADE4, PPAT*, shows only partial complementation. We have used several strategies to understand this phenotype, that point to *PPAT/ADE4* as the central regulatory node. Pathway metabolites are responsible for regulating *PPAT’s* protein abundance through transcription and proteolysis as well as its enzymatic activity by allosteric regulation in these yeast cells. Extensive phylogenetic analysis of PPATs from diverse organisms hints at adaptations of the enzyme-level regulation to the metabolite levels in the organism. Finally, we isolated specific mutations in PPAT as well as in other genes involved in the purine metabolic network that alleviate incomplete complementation by *PPAT* and provide further insight into the complex regulation of this critical metabolic pathway.

## Introduction

Since the late 80’s scientists have attempted to humanize yeast genes to study fundamentally conserved biological processes. However, due to lowered DNA sequencing and synthesis costs and improvements in molecular biology methods, recently there has been an increasing interest in human to yeast gene exchange (Kachroo et al., 2015). The growing field of synthetic biology has allowed scientists to implant entire pathways e.g. for secondary metabolites into yeast using the organism as factory for producing a desired compound (Borodina and Nielsen, 2014). However, with quite a few very extensive studies aimed at swapping single genes from human to yeast (Brachmann et al., 1996; Hamza et al., 2015; Kachroo et al., 2015) and also several attempts at pathway swapping (Choi et al., 2003; Kuijpers et al., 2016), full pathway transplantation between human and yeast has been elusive. We chose the adenine de novo pathway as a candidate for a full pathway transplant of a highly conserved metabolic pathway. The genes involved in this pathway and the regulation thereof have been well-studied in yeast cells; several of the genes in the pathway have been previously shown to be individually complemented by the human orthologs (Schild et al., 1990).

The architecture and regulation of the native de novo biosynthetic pathway have been extensively studied in *S. cerevisiae* and were reviewed by (Ljungdahl and Daignan-Fornier, 2012) (Figure 1). Purine biogenesis, a metabolically costly pathway, is regulated by a few critical metabolites. PPRP is a metabolite common to many pathways and enters the de novo pathway at its first committed step, catalyzed by Ade4/Ppat which represents a key regulatory node. AICAR and SAICAR intermediates (see below) are feed-forward transcriptional inducers of the remainder of the pathway (Rebora et al., 2001). Pathway products, such as AMP and GMP, allosterically inhibit PRPP binding to Ade4 when in high levels (Rebora et al., 2001), a finding that was also shown to be true in bacteria (Smith, 1998). In this report, pathway swapping has revealed an additional layer of regulation mediated by nucleotides that induce changes in protein abundance, probably through degradation. All of these together and an extensive phylogenetic analysis, revealing the importance of an organism specific “metabolic setpoint” for the regulation of the purine biosynthesis pathway.

**Figure 1:**
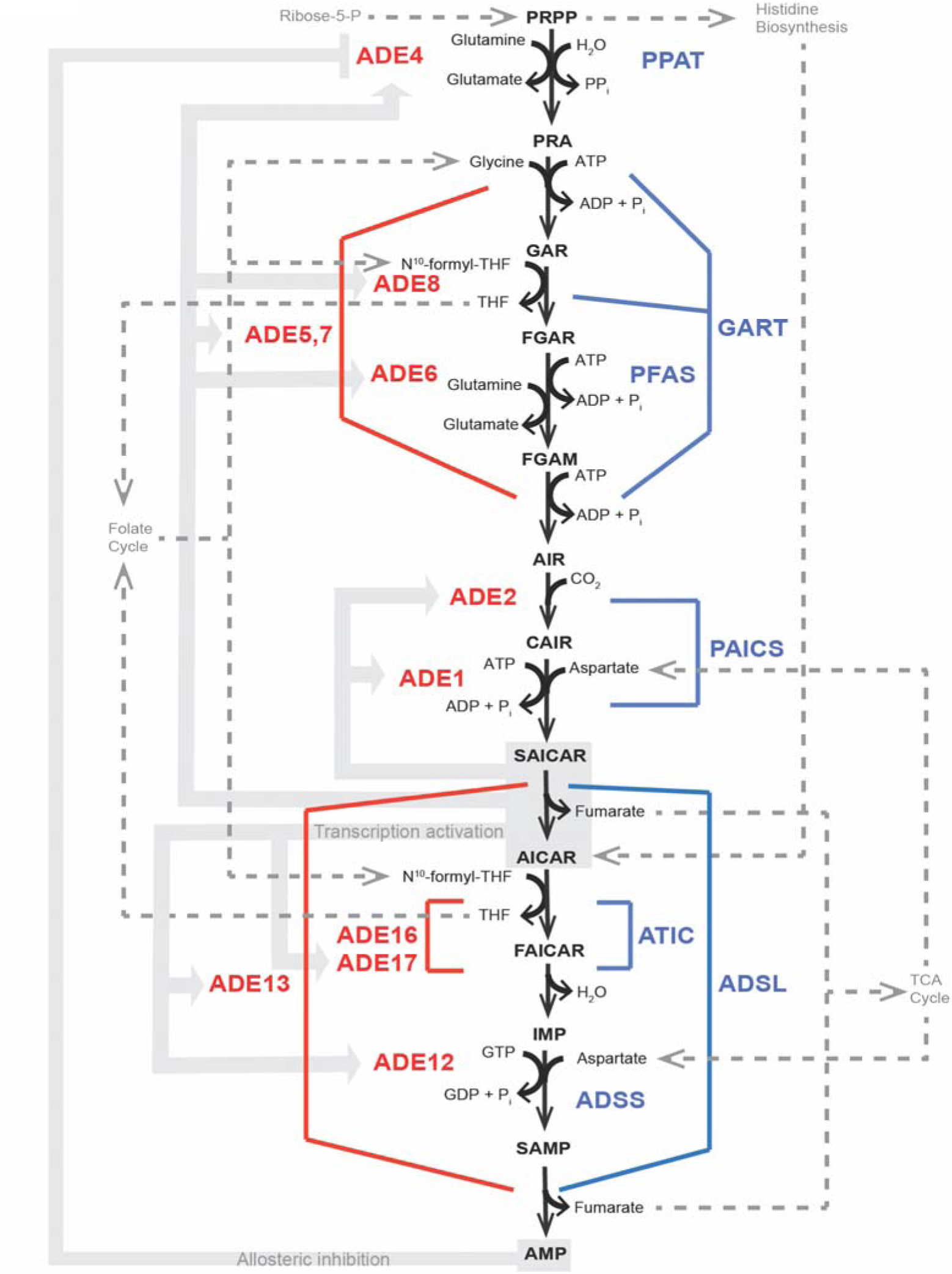
Yeast and Human Adenine de novo pathway. Schematic representation of the adenine de-novo biosynthesis pathway. Red, yeast genes; blue human. Grey dotted lines represent different metabolic pathways that interact with the adenine de-novo pathway through its metabolites. Background grey arrows indicate previously known transcriptional and allosteric regulation.

## Results

### Construction and testing of a neochromosome

In order to fully transplant a functional human pathway into a *Saccharomyces cerevisiae* cell, expression of multiple genes, each individually transcribed in the appropriate context and expression level is likely required. To accomplish these goals we expressed the adenine de-novo pathway human structural genes, recoded to yeast optimality, controlled by orthologous regulatory elements (Promoter and terminator regions). The human adenine *de-novo* pathway contains 12 steps, catalyzed by 7 distinct proteins, some of which harbor more than one enzymatic activity (Figure 1). We used a combination of yeast golden gate (yGG) assembly (Agmon et al., 2015) to build individual transcription units (TUs) followed by versatile genetic assembly system (VEGAS)(Mitchell et al., 2015) to construct a full length neochromosome in two consecutive steps (see methods).

PRO (promoter) and TER (terminator) parts from the yeast genome were employed as regulatory elements. Human CDSs were codon optimized for expression in *S. cerevisiae* and yGG compatible(Agmon et al., 2015). TUs were constructed with VEGAS adaptors (VA) as described (Mitchell et al., 2015) (Figure 2A). The correct final structure of the neochromosome (Figure 2B) was verified by DNA sequencing.

**Figure 2:**
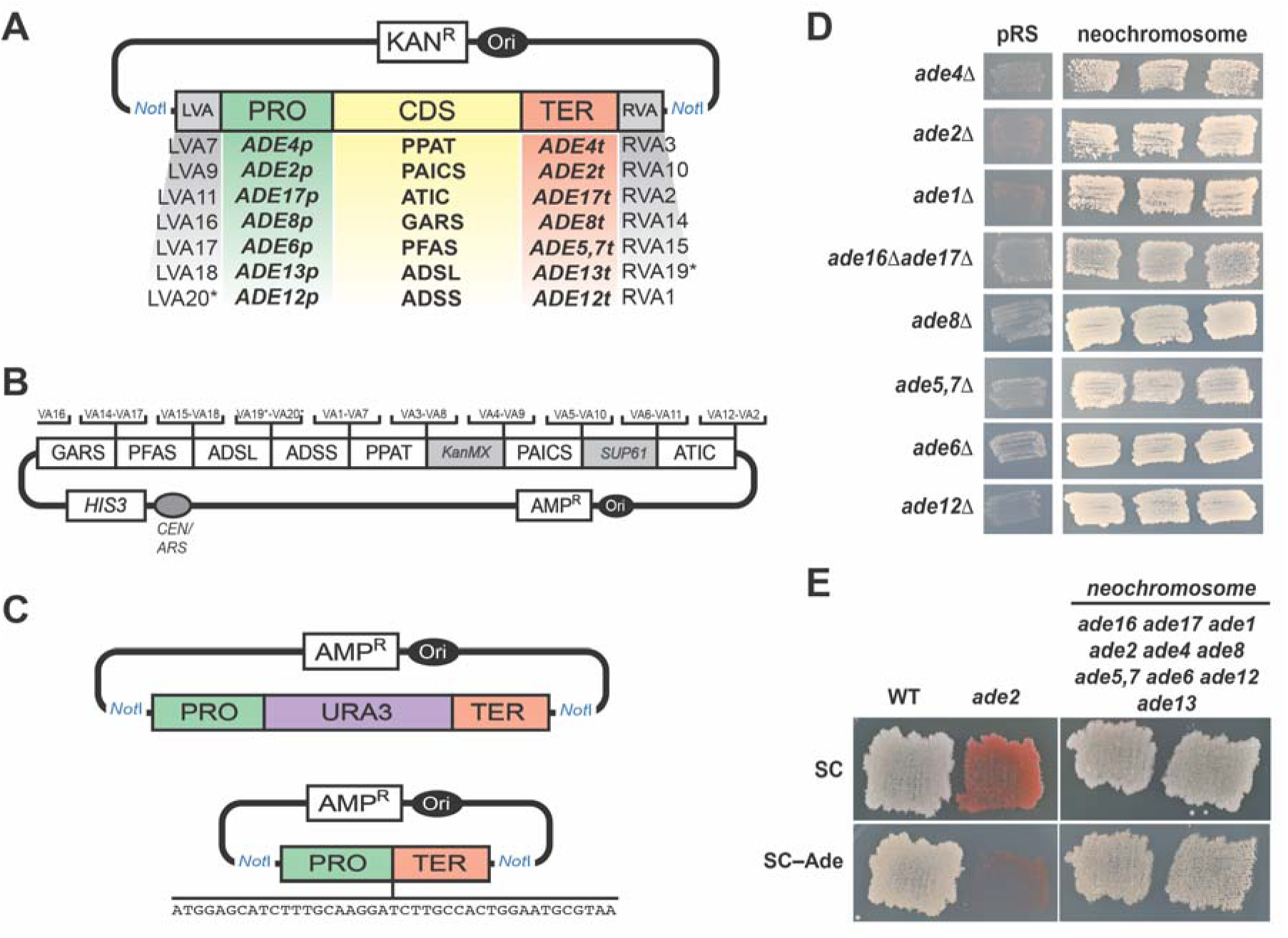
Schematic representation of neochromosome assembly and humanized yeast strains construction. **(A)** Using yeast golden-gate assembly(Agmon et al., 2015; Mitchell et al., 2015) we cloned each human gene (synthesized codon optimized for expression in *S. cerevisiae*) with its yeast ortholog’s promoter and terminator and appropriate adaptors for VEGAS assembly (VA1-VA20)(Mitchell et al., 2015) of the neochromosome (asterisk indicate previously unpublished VEGAS adaptors, for details see methods section). **(B)** Complete design of the human Purine de-novo Neochromosome for expression in yeast cells. Neochromosome was cloned in yeast cells in two steps, first step was into a pRS415 [*LEU2*] backbone and included, *PPAT, KanMX*, PAICS, *SUP61* and ATIC. The second step included overwriting (“e-SwAP-In”) of the *LEU2* marker with a *HIS3* marker and adding, GARS, PFAS, ADSL and ADSS to the neochromosome. **(C)** In order to delete the yeast *ADE* genes, we used yGG to assemble two constructs for each gene: a *URA3* integrating cassette (left) flanked with the gene’s promoter and terminator and a Linker cassette (right) also flanked with the gene’s promoter and terminator. For each gene the *URA3* cassette was first integrated, followed by erasure of the *URA3* gene by the Linker cassette. **(D)** Single gene complementation by the adenine de novo neochromosome. Yeast cells carrying single gene deletions in the yeast genes involved in the adenine de novo pathway were transformed with either the neochromosome or an empty vector with the same selection marker. Three individual colonies from the transformation were assayed for their ability to grow on media without adenine. For *ade16* and *ade17*, adenine auxotrophy is only observed in a double mutant. **(E)** Comparing growth on media with and without adenine shows complementation in the humanized strain deleted for all yeast genes and expressing all human genes from the neochromosome. Comparison is to a *ade2*Δ strain that cannot grow on media without adenine and a wt (BY4741) that can grow on both.

In parallel to assembling the humanized adenine *de-novo* pathway it is necessary to delete the yeast genes involved in the pathway in order to prove that the full heterologous pathway is functional. The 12-step yeast adenine *de-novo* pathway is catalyzed by 10 enzymes (Figure 1). In order to delete all 10 genes, using yGG assembly, we constructed two deletion plasmids for each gene, using the same flanking regions used as regulatory elements to express the human genes (Figure 2C). One plasmid insert consisted of a *URA3* gene flanked by each target gene’s upstream and downstream flanking regions, and was used to delete the target ORF via single step gene replacement. The second plasmid insert consisted of the same flanking sequences separated by a linker sequence, this insert fragment was used subsequently to delete each *URA3* gene using 5-FOA counter-selection (Boeke et al., 1987). This methodology allowed sequential deletion of multiple target genes without employing multiple markers. This exercise was performed in both *MAT**a*** and *MAT**α*** haploid strains.

The neochromosome was transformed into the multi-deletion strain as well as each individual adenine auxotrophic mutant available, and growth in the absence of adenine was assessed (Figure 2D and Figure 2E). The humanized strain grows in the absence of adenine, demonstrating the transfer of the ability to synthesize purines de novo in the complete absence of adenine supplemented from the medium. However, growth on this medium, but not on medium containing adenine, was substantially slower than the wild type.

### Partial complementation of *ade4*Δ by human PPAT

We further examined the extent of complementation by examining growth rates in adenine-free medium compared to a wild-type (WT) yeast strain. The fully humanized strain showed a significantly longer doubling time (530 min, compared to 194 min for WT). Since the yeast genes were deleted sequentially, we mapped the defect(s) by transforming the neochromosome into the multi-deletion strain series. Examination of doubling times revealed that the most significant effect on growth resulted from deletion of *ade4* (Figure 3A). All strains grew equally well in the presence of adenine, suggesting that slow growth reflects a partial failure to complement. *ADE4* encodes phosphoribosylpyrophosphate amidotransferase; like its mammalian and bacterial counterparts *PPAT* and *PurF*, it uses PRPP (phosphoribosyl pyrophosphate) as a substrate.

**Figure 3:**
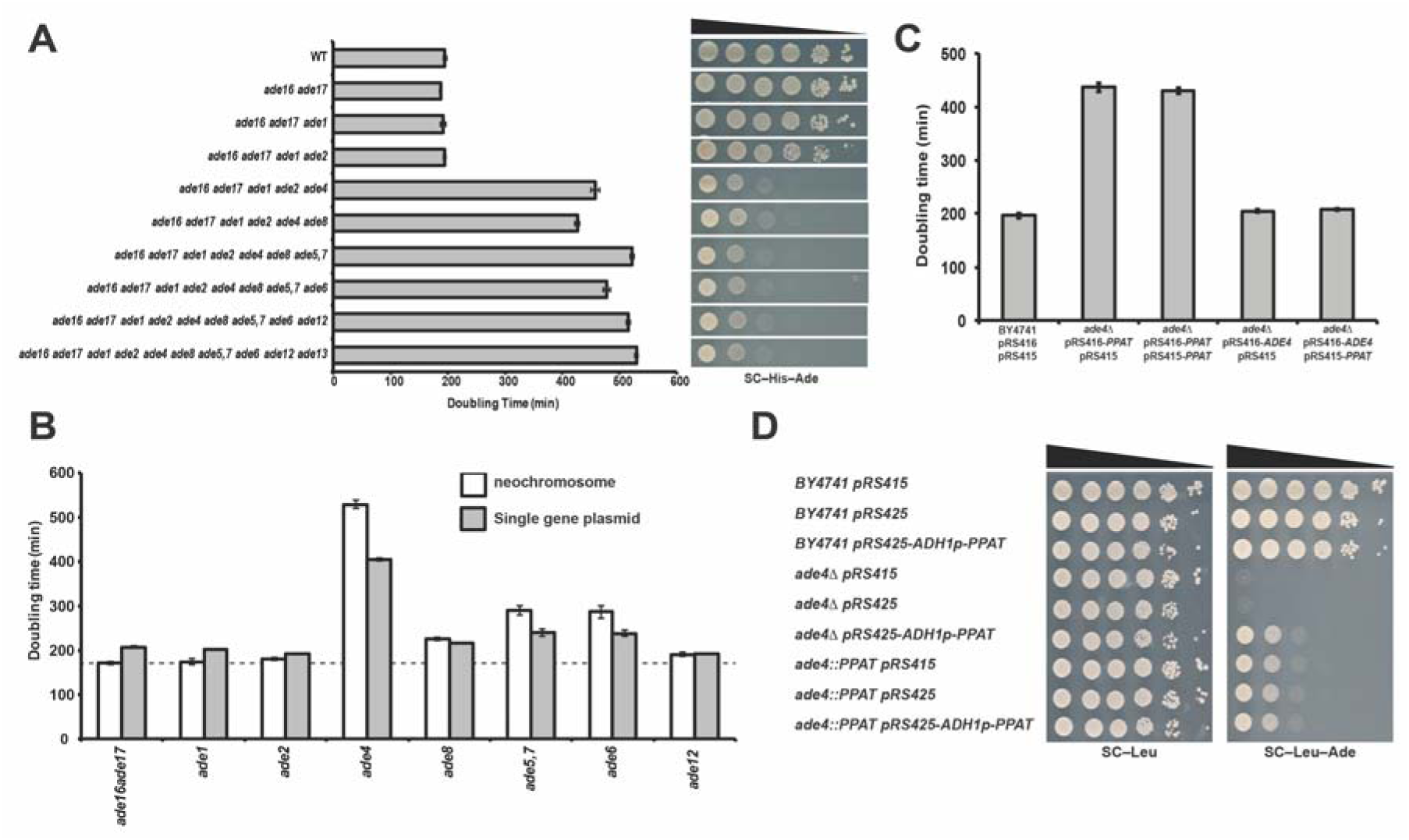
Partial complementation of ADE4 by its human ortholog PPAT. **(A)** Graphic representation of doubling time in medium without adenine of strains with increasing number of genes deleted from the de-novo pathway indicates a dramatic increase in doubling time following *ADE4* deletion (all strains carry the neochromosome). Dot assay on the left shows a similar trend to the growth assay, showing a dramatic growth defect following deletion of *ADE4*. **(B)** Graphic representation of doubling time in medium without adenine. We verified incomplete complementation of *ade4* by its human homolog *PPAT* by testing single gene deletions and comparing neochromosome complementation and single gene plasmids in medium without adenine. This shows that similarly to B, the most dramatic increase in doubling time results from *ade4::PPAT*. Dotted line represent a wild-type strain grown in the same medium. Right, dot assay showing drop in growth rate on medium without adenine. **(C)** Graphic representation of doubling time in medium without adenine of wt (BY4741) and *ade4*Δ strains carrying: pRS416-*PPAT* and pRS415, pRS416-*PPAT* and pRS415-*PPAT*, pRS416*-ADE4* and pRS415 or pRS416*-ADE4* and pRS415-*PRAT*. Doubling time was high with both one or two copies of the PPAT gene, only in those strains carrying *ADE4* plasmid was the doubling time similar to wild type. Thus, multiple copies of PPAT do not restore growth without adenine. **(D)** A dot assay examining the effect of overexpression of *PPAT*. We transformed wt, *ade4*Δ and *ade4::PPAT* strains with high copy number plasmid carrying the PPAT gene under the expression of a strong *ADH1* promoter. Increasing expression of *PPAT* did not have an effect of growth without adenine.

To verify that complementation by *PPAT* was responsible for the growth defect, the complementation ability of each human gene was tested in individual deletion mutants (Figure 3B). In agreement with the results of the multi-deletion strains, *PPAT* showed only partial complementation in an *ade4* deletion background. Evaluation of strains carrying either one or two copies of *PPAT* revealed no impact of the extra copy on doubling time (Figure 3C). In contrast, providing a single copy *ADE4* plasmid led to growth similar to the wild-type strain (with or without a *PPAT* plasmid). A second attempt at increasing expression was to construct a high copy number plasmid (2 micron) to express *PPAT* under a strong and constitutive *ADH1* promoter. We transformed this construct to BY4741, *ade4*Δ and *ade4::PPAT* (*ADE4* ORF replaced by *PPAT*). Growth without adenine of cells lacking native *ADE4*, and expressing PPAT, either from the genome, a plasmid or both, was similar (Figure 3D). These results suggest that the observed partial complementation phenotype is not simply due to *PPATs* mRNA expression level. To further investigate the differences between Ade4 and Ppat proteins, a V5-tagging strategy was used. Interestingly, whereas Ade4 could only be tagged at its C-terminus, Ppat could only be tagged at its N-terminus whilst preserving enzymatic activity (Figure S1A-B). Protein levels were examined using immunoblot analysis in media with or without adenine for 1, 3 and 6 hrs (Figure 4A). These results indicate much lower levels of Ppat protein compared to Ade4. In addition, even when Ade4 and Ppat expression was induced from an estradiol inducible promoter, the level of Ade4 was much higher than Ppat (Figure S1C).

**Figure 4:**
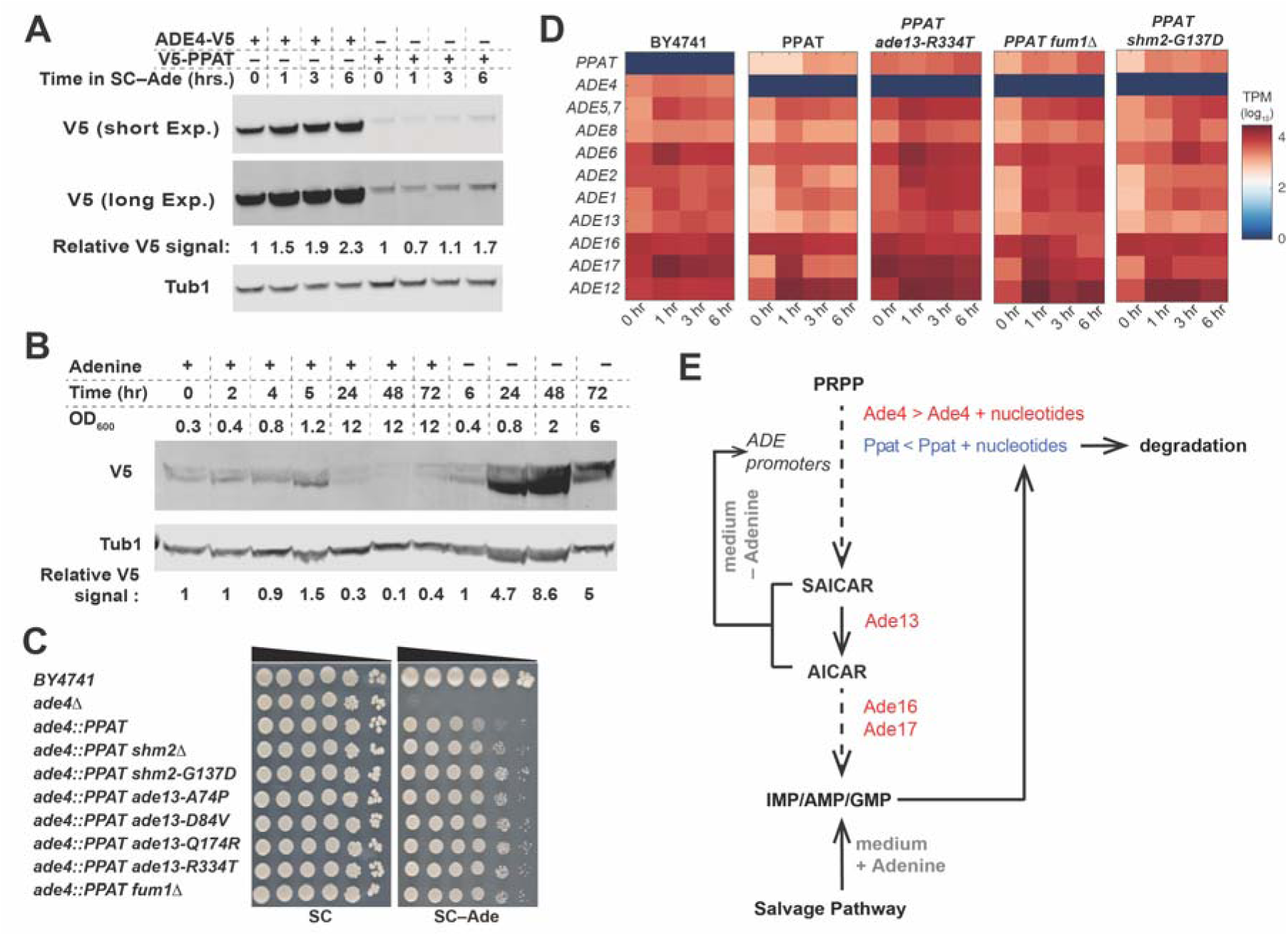
Investigating incomplete complementation of PPAT. **(A)** Immunoblot of cells expressing either Ade4 tagged with V5 or Ppat tagged with V5 integrated into the native *ADE4* locus and transcribed from its native promoter. Following over-night growth in media with adenine and transferred to media without adenine for 1, 3 and 6 h. Ade4-V5 cells show a much higher level of expression of the protein compared to V5-Ppat expressing cells. In both there is an increase in protein level during prolonged time in media without adenine. Relative V5 signal is calculated V5/Tub1 divided by the signal in time 0 h. **(B)** Immunoblot of cells expressing Ppat tagged with V5 in media with or without adenine. Ade4Δ cells were transformed with a CEN/ARS plasmid (pRS415) expressing wither V5-Ppat under an estradiol inducible promoter^15^, for details see methods. Following over-night growth in media with adenine and l μM estradiol, cells were diluted to 0.3OD and split into media with estradiol with or without adenine. Samples were taking in different OD_600_ and time points as indicated. Quantification of the amount of Ppat (relative V5 signal is calculated V5/Tub1 divided by the signal in time 0 h.) shows accumulation of significantly more Ppat in media without adenine. Results were repeated in two biological replicates. **(C)** Reconstitution of mutations found in suppressor strain genomes. Deletion of *SHM2* alleviates the PPAT phenotype similarly to the G137D mutation, indicating that the latter is probably a loss of function mutation. The *ade13* mutants show a mild increase in growth of the *ade4::PPAT* strain on SC–Ade. *fum1* deletion shows increase growth of PPAT strain in SC–Ade, indicating that the multiple mutants found in our suppressor screen are probably loss of function mutations. **(D)** Heat-map showing *ADE* gene transcripts in the yeast strains tested (BY4741, *ade4::PPAT, ade4::PPAT ade13-R334T, ade4::PPAT fum1*Δ and *ade4::PPAT shm2-G137D*) grown in SC and moved to SC–Ade. Sampled were collected after 1, 3 and 6 h for RNA preparation. RNA-seq analysis was performed on all samples and results are presented in a heatmap (scale represents log_10_, transcripts per million [tpm]). **(E)** Schematic representation of our proposed model for the complex regulation of the de-novo purine pathway through PPAT. In media with adenine salvage pathway is responsible for IMP/AMP/GMP synthesis. Nucleotides bind to Ade4 and allosterically inhibit PRPP binding, while maintaining basal level of active Ade4 (>). In Ppat expressing cells, nucleotides bind to Ppat and cause its degradation resulting in a very low level of active Ppat (<). When moved to media without adenine, in *ADE4* cells, basal level of Ade4, and the other *ADE* genes, start the de-novo pathway and increase SAICAT/AICAR levels that then through a positive feedback loop cause the overexpression of most *ADE* genes. In *PPAT* cells, due to the low basal level of *PPAT* and *ADE* genes, feedback loop is inhibited and only after nucleotides levels drop significantly active Ppat can accumulate and start the de-novo pathway. See also Figure S1 and Figure S2.

Studies in yeast and other organisms have shown that there is regulation via both allosteric and transcriptional control of the purine de-novo pathway (Daignan-Fornier and Pinson, 2012; Smith, 1998). Negative feedback inhibition by the products of the pathway occurs through allosteric inhibition of the *PPAT/ADE4/PurF* active site. In addition, in vitro tests of the activity of PurF from *E. coli* and its *B. subtilis* ortholog, revealed inhibition by distinct nucleotides(Smith, 1998). A distinct type of transcriptional regulation revealed in yeast is represented by a positive feedback loop based on binding of two pathway intermediates, SAICAR (Phosphoribosyl-aminoimidazolesuccinocarboxamide) and AICAR (aminoimidazolesuccinocarboxamide), to transcription factors that induce transcription of *ADE* genes (Rebora et al., 2001). For this dual regulation to work, a basal level of all Ade proteins must in the cells grown with adenine; this basal level is then induced on adenine depletion. In addition, previous studies have shown that binding of nucleotides to Ppat causes a conformational change (Holmes et al., 1973a; Holmes et al., 1973b; Smith, 1998). Thus, we hypothesized that PPAT might be sensitive to a certain level of nucleotide/purines in the yeast cells which inhibits it and possibly targets it for degradation. Such post-translational control is consistent with the observed lack of increased complementation with overexpression/increased copy number. This would result in a basal level too low to enable an effective response to subsequent purine depletion. A predicted secondary consequence of this would be a similarly reduced basal level of the other Ade gene mRNAs, due to a low level of the feed-forward inducers.

To test this hypothesis we examined the effect of adenine starvation on both mRNA and protein levels. We constructed a system in which a V5 tagged version of Ppat is expressed from an estradiol inducible promoter (for details see (Agmon et al., 2017)). We grew the cells in +adenine +estradiol medium for 24 hrs followed by dilution into either +Ade +estradiol or −Ade +estradiol medium. As shown in Figure 4B, there is significantly more Ppat in the cells grown in medium lacking adenine. This suggests a mechanism by which Ppat protein is sensitive to intracellular nucleotide levels available in the adenine-replete medium-grown yeast and that in the presence of the normal concentration of intracellular nucleotides/purines found in those cells, Ppat is rendered less stable, suggesting post-translational regulation via degradation of enzyme that is allosterically inhibited by nucleotide binding (Figure 4B).

We also compared the metabolome and transcriptome of the *ADE4* (BY4741) yeast cells and *ade4::PPAT* cells (*PPAT* strain) in media with or without adenine. For metabolomics analysis, we constructed otherwise prototrophic strains of both our WT strain (BY4741) as well as the *ade4::PPAT* strain, which show identical results to an auxotrophic strain (BY4741 auxotrophies) when grown in SC or SC–Ade media (Figure S1D). For both metabolome (Table S1) and transcriptome analyses, cells were transferred from SC to SC–Ade medium and samples were collected at 0, 1, 3 and 6 hrs. (see Methods). The metabolome and transcriptome results revealed several very interesting observations. 1) For most purine nucleotides tested (except GMP), their level paradoxically drops once PPAT cells are moved to SC–Ade; levels do not recover even after 6 hrs, whereas in the WT the levels are as expected higher than in SC (Figure S1E). The PPAT cells thus have a problem adapting to adenine depletion. Supporting the idea of post-transcriptional regulation of PPAT, the transcriptome of PPAT cells show much lower levels of most of the other *ADE* gene transcripts (Figure 4D), even in adenine replete medium. 2) AICAr (non phosphorylated AICAR) levels gradually increase in WT cells, whereas in PPAT cells there is a rapid increase at 1 hr followed by a decrease at 3 hrs and another increase at 6 hrs. AICAR metabolism to IMP is catalyzed by Ade16 or Ade17 (Figure 1). Although these two genes are redundant, their expression pattern is distinct; *ADE17* is induced in response to purine depletion whereas *ADE16* expression is constitutive (Figure 4D and (Denis et al., 1998; Tibbetts and Appling, 2000)). If we assume that AICAr level can serve as a proxy for AICAR level, examination of the expression level of *ADE17* might explain AICAr behavior in *PPAT* cells. In *PPAT* cells grown in SC medium *ADE17* expression is very low (Figure 4D), moving the cells into SC–Ade medium causes an accumulation of AICAr until Ade17 level is high enough to process AICAR. Only after 3 hrs, the level of AICAr drops causing the level of *ADE17* expression to drop and thus showing accumulation of AICAr again at 6 hrs 3) The level of PRPP drops in PPAT cells in SC–Ade media (Figure S1E). PRPP is the substrate of PPAT (Figure 1), and its binding to the active site is competitively inhibited by nucleotides (Smith, 1998). However, the formation of PRPP requires P_i_, the level of which is regulated by the *PHO* genes that are also induced through SAICAR/AICAR (Pinson et al., 2009). Thus, if PRPP levels are also tightly linked to *PPAT* expression, it makes PRPP levels an additional path for *PPAT* activity regulation. Finally, in contrast to AMP and IMP, GMP levels do not drop rapidly in *PPAT* cells (Figure S1E). This makes GMP a potential candidate for a primary inhibitor of Ppat.

### Suppressor analysis

We isolated a large number of independent spontaneous suppressors of the *PPAT* partial complementation phenotype in both *MAT**a*** and *MATalpha* strains by mass-transferring and growing hundreds of independent populations of cells in adenine-free medium. We then picked the strongest suppressors (Figure S1F) from each mating type and performed whole genome sequencing (WGS) on 36 of them. The mutations, none of which mapped to PPAT, identified in each suppressor are listed in Table S2. Previous work showed that adenine starvation is highly mutagenic in yeast (Achilli et al., 2004), potentially accounting for the high number of mutations found in some suppressor strains. Recurrent mutations were subsequently introduced individually into a clean *ade4::PPAT* strain to identify those sufficient to confer enhanced growth on adenine free medium. Consistent with our hypothesis, multiple recurrent mutations were mapped to genes with roles affecting SAICAR and AICAR levels. *ADE13* encodes Adenylosuccinate lyase and is a “conditionally essential” gene that catalyzes two steps in the adenine *de-novo* pathway. While it cannot be deleted from WT due to accumulation of the toxic intermediate SAICAR, it can be deleted if any upstream step is blocked. Four *ade13* mutations arose in the suppressor strain collection (A74P, D84V, Q174R and R334T). These missense mutations lie near but not in putative catalytic/active site surfaces (Ray et al., 2013). We reconstituted the four mutations in the *ade4::PPAT* strain and all four partially alleviated the slow growth phenotype (2G4, 2D1, 2F2 and 1B5 respectively, Figure 4C) and were recessive (Figure S2). In addition, we performed RNA-Seq analysis using the CEL-Seq2 protocol on PPAT ade13-R334T cells (Figure 4D). Similarly to the results obtained by Rebora and colleagues (Rebora et al., 2001) for *ade13* mutants, this suppressor was shown to have higher basal RNA levels of most *ADE* genes as well as PPAT, and also much higher induction of *ADE* genes in response to adenine deprivation. Therefore, *ade13* mutants that increase SAICAR levels increase expression of *ADE* genes generally, promoting pathway flux and improving *ade4::PPAT* strain growth on medium without adenine. Similarly, a loss of function mutation in *FUM1*, encoding fumarase, is predicted to cause accumulation of fumarate, which should shift accumulation toward SAICAR. As shown in Figure 4D, although basal level of *ADE* genes in *ade4::PPAT fum1Δ* cells are unchanged, induction of *ADE* genes after 1, 3 and 6 hrs in SC–Ade is much higher than in the control cells. Finally, recurrent mutations in *SHM2*, including a loss of function mutation, are predicted to cause accumulation of THF, a product of the Ade16 /Ade17 reaction, and to shift accumulation of AICAR precursor away from the product, FAICAR (Figure 1). Consequently, in PPAT *shm2-G137D* cells *ADE* gene expression in SC–Ade media is slightly elevated compared to the control (Figure 4C-D). These results further indicate that overexpression of PPAT alone is insufficient to fully complement *ADE4*, however, this can be mildly alleviated by also overexpressing the other enzymes in the pathway through slight increases in accumulation of the pathway metabolites that compensate by enhancing expression through the feed forward loop. A model summarizing our findings is depicted in Figure 4E.

### A zoo of PPATs

To further clarify the apparent difference between Ade4 and Ppat, we evaluated nearly 70 *PPAT* orthologs (Table S3). The codon optimized orthologs were cloned in a CEN/ARS plasmid (pRS415) with *ADE4* control sequences as described for human PPAT. We transformed the plasmids into an *ade4*Δ strain and examined their growth properties (Figure 5A and Figure S3A). A phylogenetic tree was generated based on a CLUSTAL-OMEGA multiple sequence alignment (Sievers et al., 2011). Surprisingly, some organism families seemingly clustered in groups with respect to their extent of complementation of *ade4*Δ; these groups were well-correlated phylogenetically. For example, all plants and archaea failed to complement fully, whereas all insects complemented almost completely. In bacteria, all gram-negative species tested complemented completely; whereas gram positive species showed variable complementation levels. Most mammals complemented similarly to human *PPAT*, except for platypus, which failed to complement, among the other Chordates tested all amphibians and fish complemented while reptiles and birds show variable complementation. Although there might be an underlying biological explanation for the differences among Chordates, a more likely explanation is incomplete or incorrect sequence assembly and/or annotation for the less thoroughly studied species. Those organisms in which the genome is well studied, and thus more accurately annotated, showed a very clear correlation between extent of complementation among the closely related species. Figure 5A shows the complementation from some of the highly studied organisms, and clearly shows that species with similar “life styles” show similar complementation. This result suggests that the underlying differences in complementation efficiency between orthologs might reflect the organism’s metabolic state and the relative abundance and balance of key cellular metabolites to which this critical enzyme is tuned.

**Figure 5:**
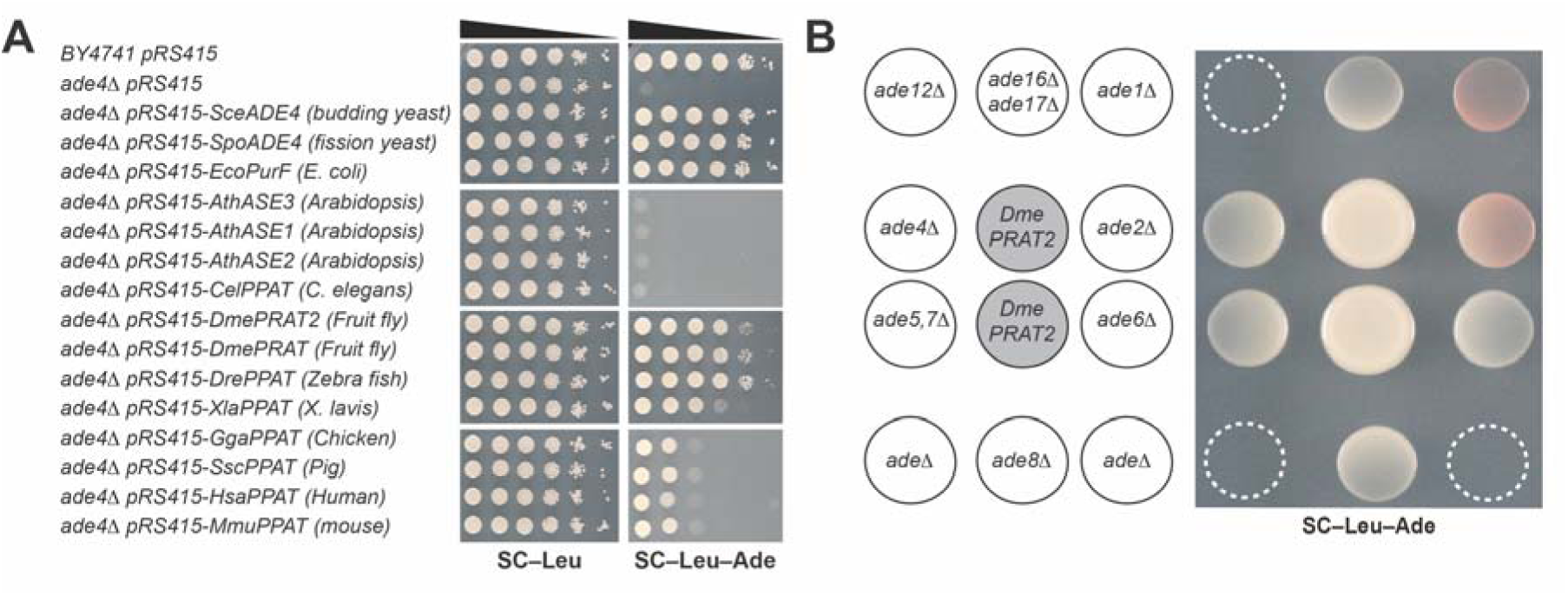
Analysis of *PPATs* from different organisms and mutations in *PPAT* PRTases domain. **(A)** Dot assay of *ade4*Δ strains carrying plasmids expressing *PPATs* from different organisms on SC/SC–Ade. Complementation is divided to 4 groups: Full complementation (Sc, Sp and Eco), nearly complete complementation (Dme, Dre and Xla), partial complementation (Gga, Ssc, Hs and Mmu) and not complementing (Ath and Cel). **(B)** Crossfeeding experiment showing that cells expressing *DmePRAT2* secret IMP, due to crossfeeding of strains deleted for different steps in the adenine de novo pathway excepting *ade12*Δ and the strain deleted for all the genes in the adenine de novo pathway (*ade*Δ). Dashed circles indicate plated cells that did not grow. See also Figure S3.

The phylogenetic analysis also unexpectedly revealed that expression of *PPATs* from a non-phylogenetically coherent subset of the organisms led to secretion of a metabolite that can cross-feed *ade4* mutants, enabling growth of neighboring cells that otherwise failed to grow on adenine-free medium (Figure S3A). To definitively identify which strains were secretors we spotted all the strains that can grow on medium lacking adenine and surrounded them with spots of *ade4*Δ cells (Figure S3B). These results show that the yeast strains expressing enzymes from *Drosophila* (both *PRAT* and *PRAT2*), *Anopheles*, purple sea urchin, shark, puffer fish, zebrafish and all three amphibians tested were “secretors”. To investigate which metabolite was secreted, we surrounded the growing spot with cells deleted for the different genes in the purine de-novo pathway, including the multi-deletion strain (*adeΔ*). The analysis suggests that overproducers apparently secrete IMP, inosine and/or hypoxanthine, and can thereby support growth for all knockout mutants in the pathway except for *ade12* (Figure 5B). Similar results have been previously observed in a dominant mutant of *Sc ade4*, named *BRA11-1*, found to secrete inosine and hypoxanthine (Rebora et al., 2001). Interestingly, this mutation was found to be located near the enzyme’s PRTase active site (Daignan-Fornier, personal communication), which binds PRPP and thus also may be the binding site for the nucleotide inhibitors of the protein (Smith, 1998). It is possible that, similar to *BRA11-1*, the ten purine secretor *PPATs* are resistant to the nucleotide levels in *S. cerevisiae* and thus do not exhibit feedback inhibition. Similarly, it may be that the cause for no or low complementation of a specific *ade4* ortholog, as in the case of human *PPAT*, is different levels of sensitivity of the specific enzyme to the nucleotide level in yeast. Thus, using mutagenic PCR we screened for mutants in PPAT that enabled better growth on SC-Ade, focusing on the PRTase domain (Figure 6 and Figure S4). Following two cycles of selection on SC–Ade, plasmid recovery and yeast transformation (see methods) faster growing colonies were observed. Plasmid recovery and sequencing revealed 5 independent mutants in the PPAT PRTase domain (Figure 6 and Figure S4). Recovered plasmids were transformed to a clean *ade4*Δ strain to check for growth phenotype, Figure 6B shows the increase in growth conferred by the mutants. We thus hypothesize that the PRTase mutants alleviate the PPAT growth defect via decreased nucleotide binding, possibly by stabilizing the protein.

**Figure 6:**
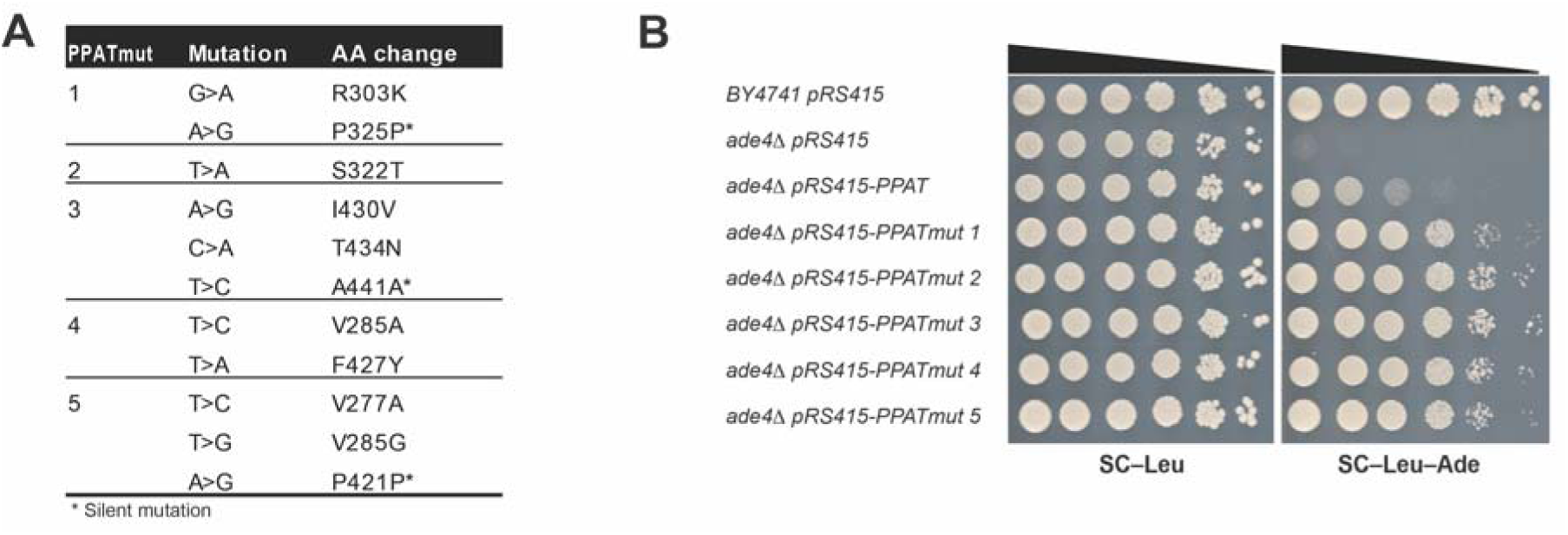
PRTase mutations partially suppress the growth phenotype of PPAT cells. **(A)** Table showing the PPAT mutants that we isolated in our PRTase mutant screen. Following mutagenic PCR and transformation into yeast, 2-3 cycles of selection (see methods) and plasmid isolation and sequencing we isolated 5 independent mutants that partially suppress the slow growth of *PPAT* expressing cells in media without adenine. **(B)** Dot assay showing the partial suppression of the slow growth phenotype by the isolated *PPAT* mutations. See also Figure S4.

## Discussion

We have engineered full pathway transplantation of a metabolic pathway from human to yeast, and shown that the human adenine de novo pathway can complement almost completely for the yeast counterpart, excepting the first pathway node, which represents the rate limiting step and thus the main regulator of the pathway, *PPAT*. This lack of complementation suggests a previously unknown protein degradation-based regulatory mechanism for the pathway. This results from inhibition of protein enzymatic activity, perhaps due to its enhanced sensitivity to distinct levels in native nucleotide levels in yeast. This in turn leads to a substantial reduction in the ability to express downstream genes in the pathway, greatly reducing growth without adenine.

Separately, analysis of diverse PPATs has revealed that their degree of complementation is highly correlated with phylogenetic distance, further supporting the above hypothesis. A subset of PPATs were shown to be overproducers of purines, further emphasizing the important role of this enzyme in regulating the entire pathway. Due to the essentiality of this pathway to dividing cells, understanding its regulation could potentially have implications in human disease. A few gene in the purine biosynthesis pathway have been shown to play a role in oncogenesis (Barfeld et al., 2015; Reitman et al., 2010). More specifically, PPAT and PAICS, were shown to be necessary for lung tumorigenesis as well as potential biomarkers for lung cancer patient survival (Goswami et al., 2015). Thus, in cancer therapy, understanding the regulation of the pathway through this single enzyme could potentially be used as a drug target for inhibiting a process which is required for the increased demand for nucleotides in rapidly dividing cells. Finally, over-secretion of purines is a phenotype that in human manifests as “hyperuricemia” due to the conversion of purines to uric acid. As shown in this report, even very small differences in the sequence of the enzyme, such as the case of human versus porcine PPAT, can make the difference between over-secreting purines and not.

## Material and Methods

### Strains and media

Yeast strains and the plasmids contained are listed in Table S4. All strains are derived from BY4741 (*MAT**a** leu2Δ0 met15Δ0 ura3Δ0 his3Δ1*) and BY4742 (*MATα leu2Δ0 lys2Δ0 ura3Δ0 his3Δ1*)(Brachmann et al., 1998). All Oligonucleotides and primers used in this work are available upon request. Media used were as follows. SD-based media supplemented with appropriate amino acids; fully supplemented medium containing all amino acids plus uracil and adenine is referred to as SC (Kaiser et al., 1994; Sherman, 2002). Through this report referring to media as with or without adenine or adenine depleted media, indicates SC media “dropped-out” for the appropriate supplements necessary to maintain specific constructs that are in the strains and either contains adenine (1.6mM final concentration) or without adenine. In addition, when cells are grown for prolonged periods of time in SC liquid media, adenine was supplements 5 time the regular amount (8mM final concentration), referred to as SCA. β-Estradiol was purchased from Sigma-Aldrich (St. Louis, MO), and 5-fluoroorotic acid (5-FOA) was from US Biological (Massachusetts, MA). Yeast strains were also cultured in YEPD medium(Kaiser et al., 1994; Sherman, 2002) or YEPD supplemented with 200 μg/ml Geneticin (G418) sulfate (Santa Cruz Biotechnology, SC-29065B)

*Escherichia coli* was grown in Luria Broth (LB) media. To select strains with drug-resistant genes, carbenicillin (Sigma-Aldrich) or kanamycin (Sigma-Aldrich) were used at final concentrations of 75 μg/ml and 50 μg/ml respectively. Agar was added to 2% for preparing solid media.

### Plasmids

Plasmids used in this work are listed in Table S5. All human genes were designed to be synthesized codon optimized for expression in *S. cerevisiae* using BioPartsBuilder (<u>http://biopartsbuilder.org)</u>. CDS were designed with appropriate overhangs and restriction enzyme to allow for cloning using yGG assembly(Agmon et al., 2015). Fragments were synthesized and cloned through Gen9Bio or GenScrip. For expression in yeast, human genes were cloned into yGG acceptor vectors (Agmon et al., 2015) flanked by their yeast orthologs promoter and terminator (Figure 2).

### Engineering of PPATs from different organisms

PPATs from different organisms were designed as described above and synthesized by Gen9Bio using yGG or using SGI-DNA BioXP 3200 system cloned in yeast. Briefly, PPAT CDS’s were synthesized (SGI-DNA) flanked by 70bp of homology to and pAV115 pre-cloned with ADE4 promoter and terminator flanking an RFP cassette flanked by *Bsm*BI sites (pNA0647). Plasmid was digested with *Bsm*BI and treated with CIP alkaline phosphatase (New England Biolabs, M0290L) recovered from gel using ZymonClean gel recovery kit (Zymo Research, D4002). It was co-transformed into *ade4Δ* cells with synthesized fragment. Transformants were screened for complementation on SC–Leu–Ade medium. For non-complementing constructs positive clones were screened on SC-Leu. For sequence verification plasmids were recovered from yeast as described below and transformed into bacteria.

### CRISPR-Cas9 system

CRISPR-Cas9 system was used to to make point mutation and protein tagging. Cas9 expression plasmid was constructed by amplifying the Cas9 gene with *TEF1* promoter and *CYC1* terminator from p414-TEF1p-Cas9-CYC1t(DiCarlo et al., 2013) cloned into pAV115 (Agmon et al., 2015) using Gibson assembly (Gibson et al., 2009). gRNAs acceptor vector (pNA0304) engineered from p426-SNR52p-gRNA.CAN1.Y-SUP4t (DiCarlo et al., 2013) to substitute the existing CAN1 gRNA with a *Not*I restriction site. gRNAs were cloned into the *Not*I site using Gibson assembly (Gibson et al., 2009). For engineering yeast using the Cas9 system, cells were first transformed with the Cas9 expressing plasmid. Following a co-transformation of the gRNA carrying plasmid and a donor fragment. Clones are then verified using colony PCR with appropriate primers.

### Neochromosome engineering

Prior to assembly of a full neochromosme we have engineered a transcription unit (TU) for each of the human genes flanked by their yeast orthologs regulatory elements and a left and right VEGAS adaptors using yeast golden gate assembly (Agmon et al., 2015; Mitchell et al., 2015) (Figure 2). These were cloned into an acceptor vector (pAV10) that carries only a bacterial selection marker (Amp^R^) and NotI sites flanking the cloned TU.

Neochromosome assembly was performed in two steps. First, the *PPAT, PAICS* and *ATIC* TUs were assembled into a *LEU2* VEGAS vector (Mitchell et al., 2015) including a *KanMx* cassette and a *SUP61* cassette. The *kanMX* cassette was used to evaluate correct assembly by replica plating to YEPD supplemented with G418. The *SUP61* cassette was included to render the neochromosome essential in yeast strains lacking the single copy and essential gene tRNA-tS(CGA)C. Following transformation onto SC–Leu plates, replica plating to YEPD with G418 showed that 100% of the colonies were G418^R^ (compared to control transformations lacking either *KanMX* or *PAICS* that showed 10 times less colonies overall and either no or very few G418^R^ colonies, respectively). 10 colonies were verified by Junction spanning PCR, and all were correct. One colony was picked to purify the neochromosome that was transformed into bacteria, and sequence verified by PacBio Sequencing.

The second step was cloning *GARS, PFAS, ADSL and ADSS* using the “eSWAP-In” method to replace the *LEU2* marker with a *HIS3* marker (pNA0177). Following transformation onto SC–His plates, replica-plating to KanMX and SC–Leu plates was performed. Of 30 transformants, two were G418^R^ and did not grow on SC–Leu, indicating successful “eSWAP-In”. Following verification of assembly using both junction spanning PCR tests, as well as additional PCR tests internal to TUs, the correct plasmids were purified from the yeast, transformed into bacterial cells and sequence verified.

### Plasmid recovery from yeast

Plasmid recovery from yeast was carried out using Qiagen buffers and Zyppy plasmid Miniprep kit (Zymo Research, D4037). Briefly, pelleted yeast cells from 1 mL of saturated culture were vortexed for 10 min. in the presence of 100 μL of 0.5 mm glass beads and 250 μL of Qiagen P1 buffer (Qiagen, 19051) supplemented with 100 μg/ml RNAase A (Qiagen, 19101). Followed by addition of 250 μL P2 buffer (Qiagen, 19052), mix and additioon of 350 μL buffer N3 (Qiagen, 19052) Following centrifugation for 10 min. supernatant was loaded on a Zyppy plasmid Miniprep kit column and washes were performed according to the manufacturer’s instruction. Plasmid DNA was eluted using sterile water heated to 55°C. 10 μL of plasmid was transformed into competent TOP10 bacteria.

### Multideletion strain construction

We first constructed two deletion constructs for each of the genes: one with a URA3 gene the other with a Linker, both flanked with the gene’s regulatory sequences (500bp upstream and 200bp downstream- Figure 2). For each genes we did a two-step deletion, first integrating a URA3 gene, followd by a second integration swapping the URA3 with a linker selected on 5FOA counter selection (Boeke et al., 1987). Due to the fact that *ADE16* and *ADE17* are redundant in their adenine auxotrophy, we started the process by deleting them in parallel in the two different mating types. The single mutants were crossed to construct a double heterozygous diploid that was then sporulated to form the double mutants. For the following deletions we deleted the genes consecutively: *ade1, ade2, ade4, ade8, ade5,7, ade6 and ade12*. For the only essential gene in the pathway, *ade13* deletion, deleting the genes upstream in the pathway renders it nonessential, thus it can be deleted in the *ade1, ade2, ade4, ade8, ade5,7, ade6 ade12* multideletion haploid strains(Hurlimann et al., 2011).

### Isolation of PPAT suppressors

*ade4::PPAT* strains of both mating type were sequentially grown and mass-transferred in restrictive conditions (SC–Ade). First we isolated single colonies and picked 96 of each mating type to grow to saturation (48 h.) in SC medium. We then diluted the cultures by 10^-3^-fold to SC–Ade and grown for 96 h., re-diluted 10^-3^-fold into SC–Ade. This was repeated 4 more times till most cultures were saturated after 48 h. In addition, from each mating type we sampled 8 of the cultures in each cycle to follow their evolvement through the experiment. Following the 5 cycles we isolated one single colony from each culture for genomic DNA preparation.

### Genomic DNA preparation

For genomic DNA preparation we used NORGEN Fungi/Yeast genomic DNA Isolation Kit (BIOTEK CORP.; 27300) following the manufacturer’s instructions.

### Genomic sequencing of suppressors

Pair-end whole-genome sequencing was performed using an Illumina 4000 system and TruSeq preparation kits. In total, 35 samples were sequenced with 3.7M – 39.5M paired reads generated per sample. The length of each read was either 101 base pairs or 151 base pairs. Quality control was performed using FastQC version 0.11.2 software (Andrews). All of the reads in the FASTQ format were aligned to the S. cerevisiae reference genome constructed starting with the sequence for control strains (strain BY4741 BY4742 genome sequences) using Burrows–Wheeler Aligner (BWA) version 0.7.8 softwarev (Li and Durbin, 2009) with -P -M -R parameter settings. Approximately 95-99% of the reads were aligned to the corresponding reference genome. GATK version 3.2 software was used to do the preprocessing (mark duplicates and local re-alignment around indels) and variant calls with -- genotyping_mode DISCOVERY -stand_emit_conf 10 -stand_call_conf 30 parameter settings(DePristo et al., 2011; McKenna et al., 2010; Van der Auwera et al., 2013). The results were subjected to a set of post-processing filters requiring: for SNPs, (i) a minimum of 10-fold coverage per variant site, (ii) WT reads in <10% of the total reads per site, and (iii) reads supporting the variant of the control sample in <5% of the total reads per site; for Indels, (i) a minimum of 10-fold indel coverage per indel site of the treatment sample, (ii) a maximum of 5-fold indel coverage of the control sample (iii) a minimum of 90% of treatment indel within the local region using 50bps flanking regions on both directions. snpEff version 4 software(Cingolani et al., 2012) was used to annotate each variant using Toronto-2012 gene GFF file.

### RNA preparation and sequencing

Cells were grown for 24 hrs in SC media to saturation, then diluted to SC–Ade media to a concentration of 10^7^ cells/ mL. Samples were collected at 0, 1, 3 and 6 hrs. span down and flash freeze in liquid nitrogen to be stored at −80C until RNA preparation. RNA was prepared as described at (Mitchell and Boeke, 2014) from 10^7^ yeast cells by using the RNeasy Minikit (Qiagen; 74106) as per the manufacturer’s instructions. In brief, cells were lysed enzymatically, and eluted RNA was treated with DNase (Qiagen; 79254) in solution before passage over a second column and elution in water. Approximately 5ng of each sample was used for RNA amplification and library preparation using the CEL-Seq2 protocol taking only ^1^/_5_ of the amplified RNA to prepare the library. Paired end sequencing was performed on the Illumina NextSeq 500. The sequencing data was de-multiplexed using the CEL-Seq pipeline (Hashimshony et al., 2016). Mapping of the reads was done using bowtie2 version 2.2.6 (Langmead et al., 2009) in the following way: for WT samples the reads were mapped using the genome of BY4741, for the mutant samples the reads were mapped using the BY4741 genome adding the human gene, PPAT. Read counting was performed using an adaptation of HTseq to count each UMI only once (Hashimshony et al., 2016). The counts were normalized by dividing the total number of mapped reads for each sample and multiplying by million [Transcript per million (TPM)].

### Metabolomic analysis

Cells were switched from SCD + 20 mg/L adenine to SCD – adenine for 1, 3, 6 hours then extracted with 75% ethanol. Supernatant (1 ml) was collected, vacuum dried, and stored at -80°C. The metabolite analysis was performed by LC-MS/MS on a Shimadzu Prominence LC20/SIL-20AC HPLC coupled to a ABSCIEX 3200 QTRAP triple quadrupole mass spectrometer as described previously (Laxman et al., 2014). Chromatographic separation was performed using a C18-based column with polar embedded groups (Synergi Fusion, 150 × 2.0 mm 4 μ, Phenomenex). Infusion quantitative optimization was performed to acquire optimal product ion mass for each metabolite. Multiple reaction monitoring (MRM) was used to detect and quantitate metabolites. The two most abundant daughter ions were used when possible and metabolite peak area was normalized to total ion content. Buffers for positive-mode analysis were formic acid method (buffer A: 99.9% H_2_O/0.1% formic acid and buffer B: 99.9% methanol/0.1% formic acid), and ammonium acetate method (buffer A: 5 mM ammonium acetate in H_2_O and buffer B: 5 mM ammonium acetate in 100% methanol). TBA method (buffer A: 5 mM tributylamine (TBA) and buffer B: 100% methanol) was used for negative-mode. The area under each peak was quantitated by Analyst software, and normalized against total ion count.

### Immunoblot analysis

For protein extraction cells expressing either Ade4-V5 or V5-Ppat were grown for 24 hrs. in SC media, followed by dilution and transfer to media with or without adenine and with or without 1μM estradiol as described above. 20 OD_600_ were collected, washed once with water and flash frozen in liquid nitrogen for storage in −80°C until protein preparation. Cells were incubated for 30 min. in NaOH buffer [150mM NaOH, 2mM DTT and 1X cOmplete, EDTA free protease inhibitor Cocktail (Roche, 11873580001)] on ice for 30 min. Following 10 min. centrifugation at 4°C, cells were lysed in 150 mL of lysis buffer [20 mM HEPES, pH 7.4, 0.1% Tween 20, 2 mM MgCl2, 300 mM NaCl, 1.5mM DTT and 1X cOmplete, EDTA free protease inhibitor Cocktail] in the presence of equal volume of 0.5 mm glass beads (1:1 cell slurry:beads) by vortexing (10 min. at 4°C). Following centrifugation (10,200 rpm, 4°C, 3 min), 150 μl of clarified whole-cell extract was collected and total protein was measured using Bradford protein assay kit (BioRad, 5000006). For estradiol experiment (Figure 3A) total cell lysate was concentrated using microcon-30kDa filter unit (EMD Millipore, MRCF0R030) before measuring total lysate concentration. Samples were then mixed with 4X LDS sample buffer (Life Technologies, NP0007). Samples were heated at 70°C for 10 min and loaded onto either a 12% pre-cast Bis-Tris gel (Life Technologies, NP0342BOX) or a 10% pre-cast Bis-Tris gel (Life Technologies, WB1202BX10) and electrophoresed in 1x MOPS buffer (Life Technologies, B000102). Protein transfer was carried out using a BioRad Trans-blot Turbo Transfer system and corresponding reagents. Anti-V5 antibody was from Sigma (V7754-4MG; Rabbit) and Anti-Tub1 (Rabbit) served as a control for loading. Secondary antibodies were from LI-COR (IRDye 800CW, 926-32210; IRDye 680RD, 926-68071). Western blots were developed and quantified using the LI-COR Odyssey and Image Studio Software.

### *ade13* suppressor mutation recessivity test

In order to distinguish whether these *ade13* suppressor mutants are hypomorphs or gain of function mutations we crossed each of the re-constructed strains to a *MATalpha* strain containing *ade4::PPAT* and a WT copy of *ADE13*. We then plated the cells on media with or without adenine. Comparing the growth of the *ade4::PPAT ade13* heterozygots with *ade4::PPAT ADE13* WT homozygous on media without adenine revealed that the mutants are recessive and thus most likely hypomorphs.

### PPAT PRTase mutants screen

We preformed mutagenic PCR of the PRTase domain of PPAT with forward primer GTTGAAATCTCTAGACACAACGTTCAAACC and reverse primer CAAGTGGTCGAATTCTGGCTTGTT using GeneMorph II Randome Mutagenesis Kit (Stratagene, 200550) following the manufacturer’s instructions. Briefly, 1.7 μg of plasmid (pNA0310) was used to amplify a 665bp fragment corresponding to the PRATase domain of PPAT (Figure S9). Product was then co-transformed to *ade4Δ* yeast cells with pNA0642 digested with BstZ17I, treated with CIP alkaline phosphatase (New England Biolabs, M0290L) recovered from gel using ZymonClean gel recovery kit (Zymo Research, D4002). For each transformation reaction 10% of transformed cells were plated on SC–Leu media to calculate transformation efficiency, 10% were plated on SC–Leu–Ade to assay for growth of mutants on media without adenine and 80% were put into SC–Leu–Ade culture for selection. Following 48 h growth in culture, plasmids were prepped from yeast as described above, transformed into bacteria, grown in bacterial growth media with the appropriate antibiotics for selection, plasmid prepped and re transformed to *ade4Δ* yeast cells for another cycle of selection. This was repeated three times in 4 independent cultures (lines). 2-4 colonies were picked from each line, plasmid prepped, transformed to bacteria, plasmid prepped and sent for sequencing to find the mutations. Out of 15 plasmids sequenced we isolated 5 different mutants of *PPAT* (Figure 6). To validate that the phenotype was a result of the plasmid mutation we re-transformed sequenced plasmids to clean *ade4Δ* yeast cells.

## Acknowledgments

We thank Boeke lab members for helpful discussions. We also thank Michael Pacold and Bertrand Daignan-Fornier for helpful discussions and generous sharing of reagents and information. This work was supported by NIH grant 1R24DK082840. J.D.B. is a founder and director of Neochromosome Inc. J.D.B. serves as a scientific advisor to Modern Meadow, Inc., Recombinetics Inc., and Sample6 Inc. These arrangements are reviewed and managed by the committee on conflict of interest at NYULMC.

